# Estimating the neural spike train from an unfused tetanic signal of low threshold motor units using convolutive blind source separation

**DOI:** 10.1101/2022.10.12.511951

**Authors:** Robin Rohlén, Jonathan Lundsberg, Christian Antfolk

## Abstract

The central nervous system initiates voluntary force production by providing excitatory inputs to spinal motor neurons, each connected to a set of muscle fibres to form a motor unit. Motor units have been imaged and analysed using ultrafast ultrasound based on the separation of ultrasound images. Although this method has great potential to identify regions and trains of motor unit twitches (unfused tetanus) evoked by the spike trains, it currently has a limited motor unit identification rate. One potential explanation is that the current method neglects the temporal information in the separation process of ultrasound images, and including it could lead to significant improvement. Here, we take the first step by asking if it is possible to estimate the spike train of an unfused tetanic signal from simulated and experimental signals using convolutive blind source separation. This finding will provide a direction for ultrasound-based method improvement. In this study, we found that the estimated spike trains highly agreed with the simulated and reference spike trains. This result implies that the convolutive blind source separation of an unfused tetanic signal can be used to estimate its spike train. Although extending this approach to ultrasound images is promising, the translation remains to be investigated in future studies where spatial information is inevitable as a discriminating factor between different motor units.

## Introduction

The central nervous system initiates voluntary force production and human movement by providing excitatory inputs to spinal motor neurons. Each motor neuron connects to a set of muscle fibres by a synaptic connection to form a motor unit (MU). The gold standard for measuring characteristics of a large population of MUs is based on high-density surface electromyography (sEMG)^1^. High-density sEMG consists of a grid of electrodes that records mixed MU activities superficially from the skin (<20 mm). Then, the activity is decomposed into single MU spike trains by blind source separation (BSS) techniques, which use the large number of electrodes. This approach provides access to the neural drive of the spinal cord via the motor neurons to specific muscles^2^. Although sEMG has been used successfully for MU analysis, it is well-known that it has limited spatial selectivity and field of view.

Ultrafast ultrasound has been shown to image and analyse voluntarily activated MUs for a large field of view in the muscle (40×40 mm)^3–8^, providing spatiotemporal mechanics at a high resolution (<1 mm and >1 kHz). This technique is based on recording radio frequency signals (B-mode images) (Fig. 1A-B), calculating displacement velocity images (Fig. 1C) and separating the images into spatiotemporal components, i.e., each component is associated with a 1) spatial image and 2) a time signal (Fig. 1D). A subset of the components is putative estimates of the 1) MU territory and 2) trains of MU twitches (unfused tetanus) evoked by the spike trains^3,4,7,8^. The spike trains are estimated based on unfused tetanic signals (Fig. 1E). Although this ultrasound-based method (Fig. 1) has great potential to provide comprehensive access to the neural drive of the spinal cord to muscles for a large population of MUs simultaneously, the method currently has a limited identification rate of the active MUs^4^.

**Figure 1.**
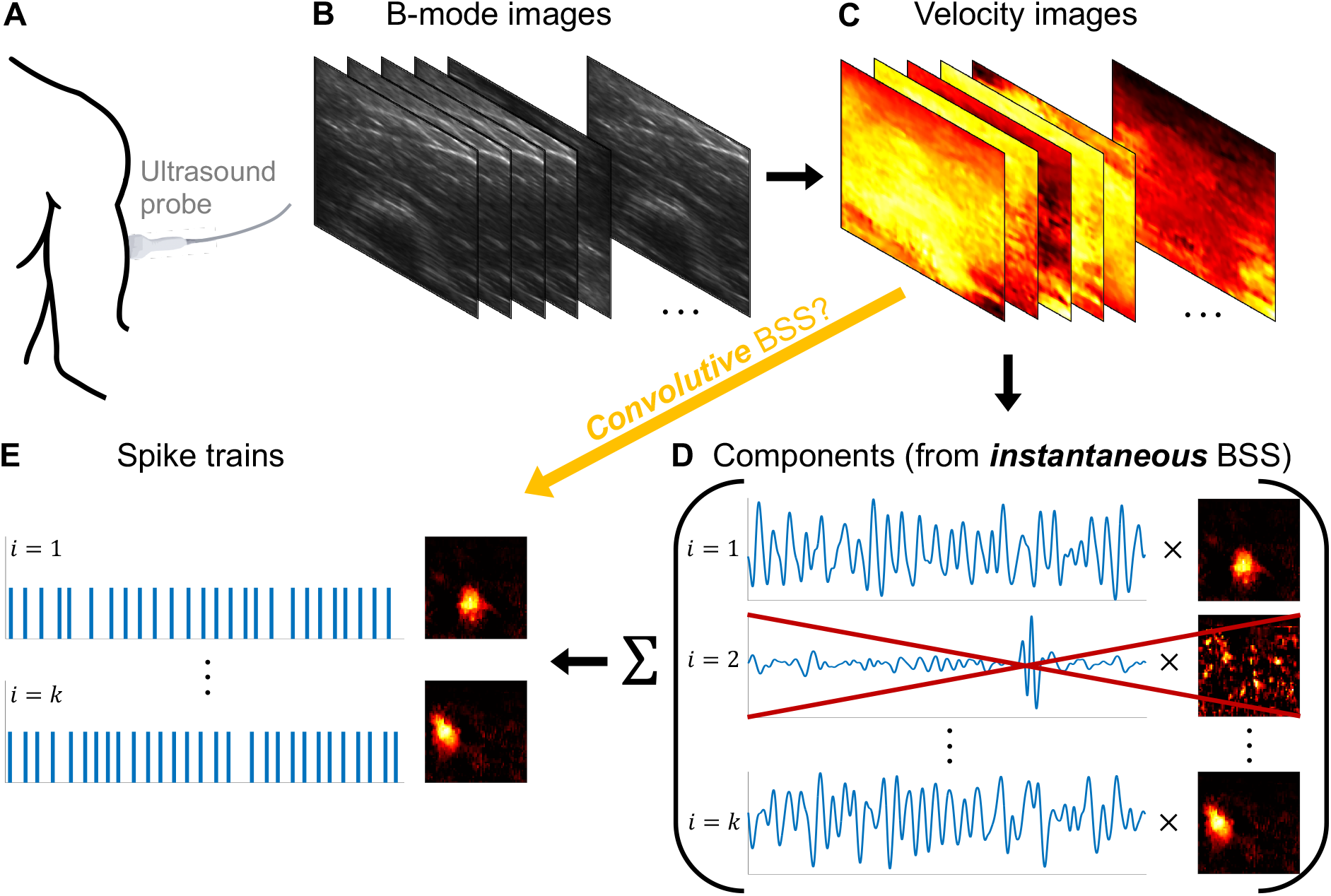
The methodological pipeline to identify voluntarily activated motor units (MUs) using ultrafast ultrasound. **A**. The ultrasound probe is placed on the skin to record data from the muscle’s cross-section (transverse view). **B**. The recorded B-mode images. **C**. Calculating displacement velocity images. **D**. Separating the velocity images into spatiotemporal components using *instantaneous* blind source separation (BSS) with focus on spatial sparsity, where each component is associated with a time signal and a spatial image. A subset of the components is putative estimates of the 1) MU territory and 2) trains of MU twitches (unfused tetanus) evoked by the spike trains. **E**. The spike trains are estimated based on unfused tetanic signals. Including the temporal *convolutive* BSS approach for the ultrasound-based decomposition method means going from velocity images directly to spike trains (from **C** to **E**).

A recent study found that methodological improvements in velocity image separation (blind source separation, BSS) should lead to a higher identification rate^6^. A methodological improvement could include more information about the MU characteristics in the separation process^6^. The reasoning is that the current method neglects the *temporal* information in the separation process and focuses on sparse *spatial* (pixel) distributions as cost functions motivated by the physical muscle unit territories (Fig. 1D). The velocity images are decomposed into all components simultaneously using a *symmetric* approach based on *instantaneous* BSS. In contrast, the state-of-the-art sEMG decomposition methods use a *deflationary* approach, i.e., estimate one component (putative MU) at a time^9–11^, and exploit the spike trains’ *temporal* sparsity through *convolutive* BSS. Including the temporal convolutive approach for the ultrasound-based decomposition method means going from velocity images directly to spike trains (i.e., Fig. 1C to Fig. 1E). This approach could lead to an improved method with a higher identification rate of active MUs.

Although including the temporal convolutive approach in ultrasound decomposition is promising, Lubel et al. (2022) found that the motion of non-MU-related structures hides a large part of the movement caused by a MU in ultrasound images^12^. This finding is in line with the motivation for “over-decomposing” with the ultrasound decomposition method^3^ (Fig. 1D), by increasing the number of components to account for non-MU-related structures. Since decomposition methods are biased toward high-amplitude signals^13^, which in this case is non-MU activity, a direct translation of the sEMG BSS algorithms to ultrasound images is not feasible. Given this, we would like to take the first step in investigating the feasibility of this translation by asking if it is possible to estimate the spike train of an unfused tetanic signal using convolutive BSS. If this is feasible, we have a clear direction for ultrasound-based method improvement through the integration of temporal information in the separation process of velocity images.

The main challenge of using convolutive BSS to estimate the spike train of an unfused tetanic signal is that the unfused tetanus of a MU will consist of *variable* successive twitches^14–16^, where the twitch duration is *longer* than the time between two succeeding spikes (i.e., an ISI). In contrast, the MU action potential is more *consistent*, and its duration is *shorter* than the ISI. Previous studies of spike train estimation of an unfused tetanic signal have exploited the similarity of each twitch rise (onset gradient) and also observed that the twitch rise duration is shorter than the ISI^17^. Assuming the twitch rise is the action potential counterpart, and the remaining twitch activity is another additive noise component, we hypothesise that estimating the spike train using convolutive BSS is possible.

This study aimed to estimate spike trains of simulated and experimental unfused tetanic signals using convolutive BSS. We used a convolutive BSS algorithm with three parameters based on a fixed-point iteration^10,11^ and peak detection^17^. We evaluated the algorithm’s parameters and its performance in terms of the estimated spike trains’ rate of agreement with 1) the ground truth for the simulations and 2) the EMG reference for the experimental data. We also explored the algorithm’s parameters’ relation with the spike delta (time difference between truth or reference spikes and the estimated spikes) and its variation.

## Methods

The overview of the methods includes 1) simulations describing the model and the parameters, 2) the experimental data, 3) the convolutive BSS algorithm to estimate the spike train, and finally, 4) the parameter and performance evaluation using simulated and experimental data.

### Simulations

#### Simulation model

We used a simulation model that generated a motor unit (MU) unfused tetanic signal *Y*(*t*) at time *t* ≥ 0:

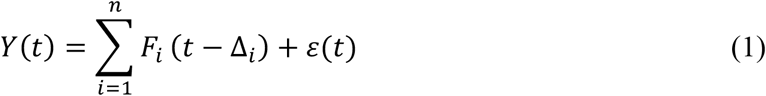

where *t* = {0,1, …, *T*} is the time and *n* is the number of times that the MU fires. *F_i_* is the *i*^th^ twitch generated by the MU at its *i*^th^ spike at time instant Δ*_i_*. The duration of *F_i_* is longer than the inter-spike interval (ISI) Δ_*i*+1_ − Δ_*i*_. *ε*(*t*) is additive noise. Note that the twitch-shape commonly varies within a contraction, which has been explained in other studies for voluntary contractions^14–16^. We express the corresponding spike train *s*(*t*) ≥ 0 as:

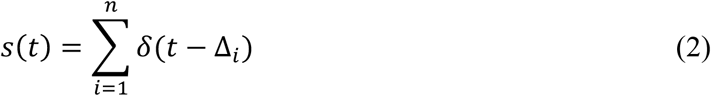

where *δ* is the Dirac Delta function.

#### Simulation parameters

Similar to a previous study^17^, each simulated unfused tetanic signal *Y*(*t*) was based on simulating *n* twitches and *n* spikes (Fig. 2). The twitches *F_i_*, *i* ∈ {1,2, …, *n*} were simulated using a twitch model^16^ with five parameters (Fig. 2A) that were randomly sampled to generate varying twitch-shapes (Fig. 2B) suitable for low threshold MUs at low force levels^18^ (see Table S1, Supplementary Material). The *n* spike times Δ = (Δ_1_, Δ_2_, …, Δ_n_) were simulated using a Gaussian renewal process such that Δ_*i*+1_ − Δ_*i*_ ~ *N*(*μ_ISI_*, CV), where *μ_ISI_* and 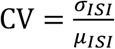 is the mean ISI and coefficient of variation^19^ (Fig. 2C). We simulated combinations of different firing rates (8, 12, and 16 Hz)^4,20^ and ISI CVs (5, 20, and 40%)^21^ to mimic the characteristics of low-threshold MUs.

**Figure 2.**
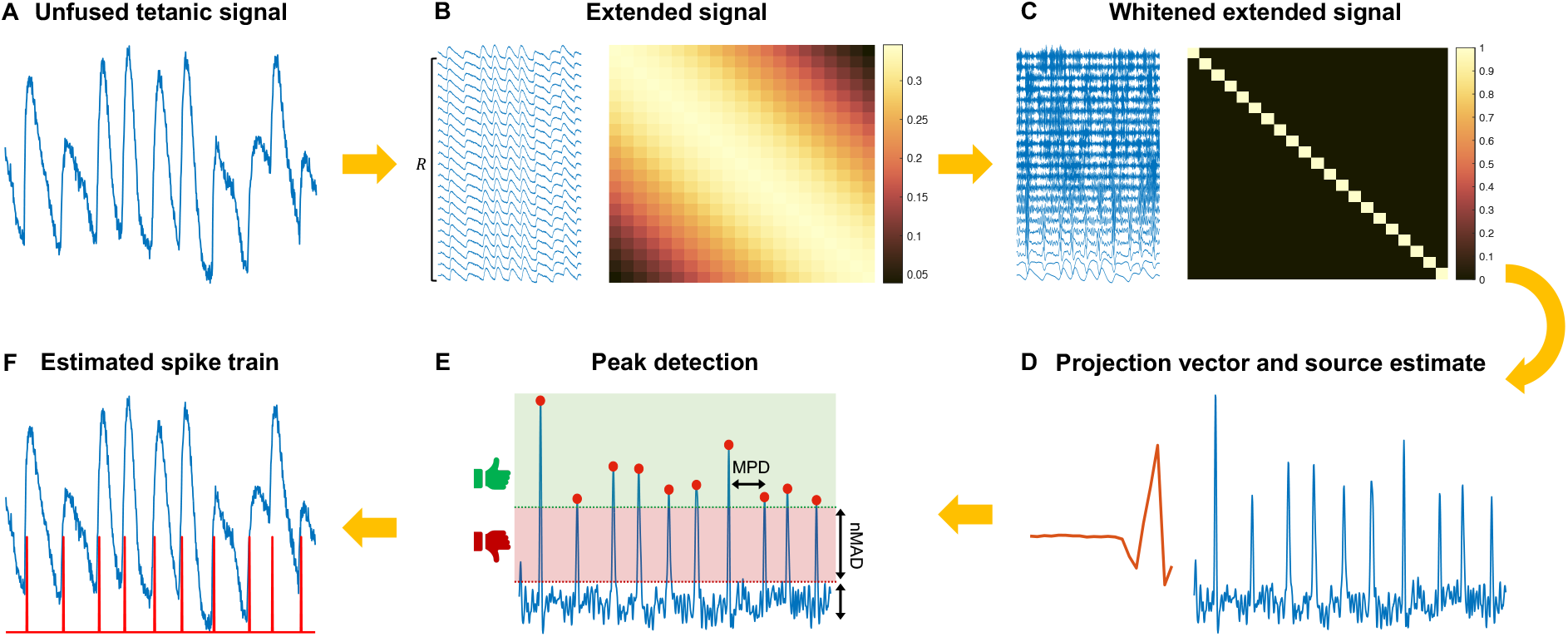
The different steps of the convolutive blind source separation algorithm. **A**. The input signal is the unfused tetanic signal. **B**. The unfused tetanic signal is extended using the extension factor *R*, where the extended signal has a dense covariance matrix. **C**. Performing whitening such that the covariance matrix is equal to the identify. **D**. The fixed-point algorithms find the separation (projection) vector (red line), and by projecting it on the whitened extended signal in **C**, the resulting output is the source estimate (blue line). **E**. The time instants of the local maxima of the source estimate were detected using peak detection with two parameters: the minimal peak distance (*MPD*) and the number of mean absolute deviations (*nMAD*). **F**. the time instants of the local maxima are the estimated firing times of the spike train ***s*** (red line).

Each twitch was summed to the respective spike resulting in an unfused tetanic signal (Fig. 2C). The unfused tetanus was *differentiated with respect to time* (from force to yank or displacement to velocity)^22^ because of the stability of its mean value, and velocity is used in MU identification using ultrasound. The last step was adding Gaussian noise so that the signal-to-noise ratio (SNR) was at a pre-defined level (10, 20, and 30 dB). For detailed information on the simulations, see Supplementary Material.

### Experimental signals

We retrospectively included, from previous studies, two datasets including 21 unfused tetanic signals^17^. The first dataset included three two-second-long signals from the biceps brachii of three healthy human subjects (28.3 ± 0.6 years; two females) at low voluntary force levels using the MU ultrasound analysis with a needle EMG spike train as reference^4^ (see Supplementary Material). The ultrasound and EMG systems were synchronized with 2 kHz and 64 kHz sampling rates. The subjects gave informed consent before the experimental procedure. The project conformed to the Declaration of Helsinki and was approved by the Swedish Ethical Review Authority (2019-01843).

The second dataset included eighteen electro-stimulated signals from functionally isolated slow MUs in the medial gastrocnemius of five adult female Wistar rats^23^. After the surgical procedure, the Achilles tendon of each rat was connected to an inductive force transducer while stretched to achieve isometric conditions where the force was measured simultaneously as a wire electrode with sampling frequencies of 1 kHz and 10 kHz, respectively. The stimulation frequencies for the first three rats were 10, 12.5, 14.3, and 16.6 Hz. Only the first three stimuli intervals were considered for the other two rats. To mimic voluntary contractions with varying stimuli intervals, the intervals between the individual stimuli were randomly set at values in the mean ISI ± 50% range. Before further analysis, the force-based signals were 1) filtered using a 6^th^ order zero-phase Butterworth bandpass filter with a high- and low-pass cut-off equal to 3 and 100 Hz and 2) differentiated. The experimental procedures followed the European Union animal care guidelines, and the principles of Polish Law on the Protection of Animals were approved by the Local Bioethics Committee. For detailed information on the surgical procedure, see Supplementary Material.

### The convolutive blind source separation algorithm to estimate the spike train

Previously a shift-invariant model has been used to describe a multichannel sEMG signal assuming 1) constant action potential waveforms for the same MU in one channel and 2) that the waveform duration is shorter than every ISI^9–11^. This study considers a single-channel unfused tetanic signal *Y*(*t*) which has the following two characteristics: 1) the twitch waveforms for the same MU vary^14–16^ and 2) the twitch waveform duration is longer than the ISI^15,24^. Given this, we express the time derivative of the unfused tetanic signal as:

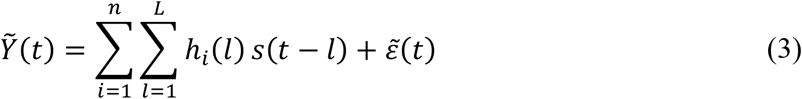

where *i* = {1, …, *n*} denote the *i*th spike, *h_i_*(*l*) is a twitch rise waveform of length *L* as a response to the *i*^th^ firing, *s*(*t*) is the spike train at time *t*, and 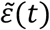 is additive noise including the remaining signal of the twitch waveforms after subtracting the twitch rise waveform. The convolutive mixture with finite impulse response filters in equation (3) can be represented as a linear and an *instantaneous* mixture by extending the observation signal, source signal, and noise using their *R* delayed versions^9–11^:

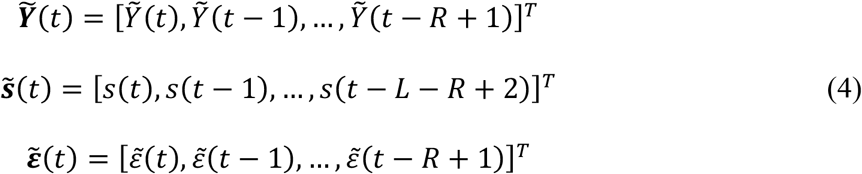

where *L* denote the twitch rise waveform length and *R* denote the extension factor. Then, the linear instantaneous mixture model is defined as:

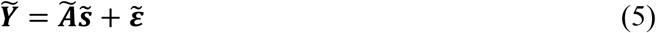

where 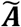 contains the twitch rise waveforms *h_i_*. To estimate the spike train ***s*** = *s*(*t*), *t* = {0,1, …, *T*} using equation (5), we used a linear instantaneous BSS model referred to as independent component analysis (ICA)^25^.

#### Algorithm

Given an unfused tetanic signal (Fig. 3A), the spike train ***s*** was estimated by solving the separation problem in equation (5) using a fixed-point algorithm^26^, which has been used for decomposing multichannel EMG signals^10,11^. After extending the signal (Fig. 3B), the next step, common to most BSS algorithms^25^, was to spatially whiten the extended observation matrix using eigenvalue decomposition (Fig. 3C). The whitened extended observation matrix had a covariance matrix equal to the identity (Fig. 3C). After whitening, the fixed-point algorithm estimates the separation (projection) vector and, thereby, the source estimate (Fig. 3D). Here, the separation vector was initialized using a gaussian random vector. The fixed-point algorithm is based on maximizing the non-Gaussian distribution of the source vector using a cost function *G*(∙)^26^. We used a contrast function to measure sparseness since the spike train is sparse (mostly zeros with a few ones). Here, we choose *G*(*x*) = log (cosh(*x*)) due to its kurtotic distribution in line with the characteristics of a spike train that is kurtotic (and skewed) in contrast to a normal distribution.

**Figure 3.**
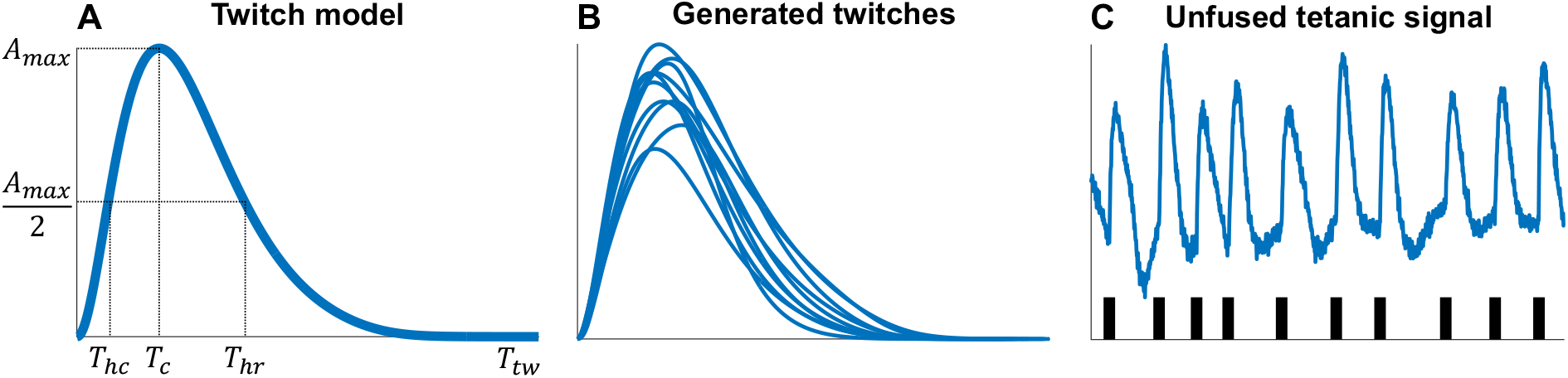
Simulating unfused tetanic signals using a twitch model and a spike train. **A**. A twitch (blue line) is generated by its five parameters (black dashed lines) of the model are the half-contraction time *T_hc_* (ms), the contraction time *T_c_* (ms), the half-relaxation time *T_hr_* (ms), the twitch duration *T_tw_* (ms), and the maximal amplitude *A_max_* (arbitrary unit, a.u.). **B**. Generated twitches using the twitch model in **A** by randomly sampling their parameters from a uniform distribution. **C**. Time instants (spikes, black vertical lines) for each twitch were simulated using a Gaussian renewal process. The unfused tetanic signal was obtained by summating the twitches in **B** at their respective time instants with added Gaussian noise to achieve a certain signal-to-noise (SNR) level. Note that the unfused tetanic signal has been differentiated with respect to time, providing unfused tetanus concerning the rate of change (e.g., yank or velocity instead of force or displacement).

After obtaining the source estimate, the spike train was estimated by peak detection to identify the time instants of the local maxima. The peak detection was based on 1) the height of the peaks and 2) the distance between consecutive peaks in milliseconds (Fig. 3E). The height of the peaks was based on the number of mean absolute distances (*nMAD*) of the source estimate. The distance between the consecutive peaks was based on the minimum peak distance (*MPD*) where the highest peak within that distance was selected. The resulting time instants of the local maxima were considered the estimated firing times of the spike train ***s*** (Fig. 3F). The overall algorithm is summarized in pseudo-code below:

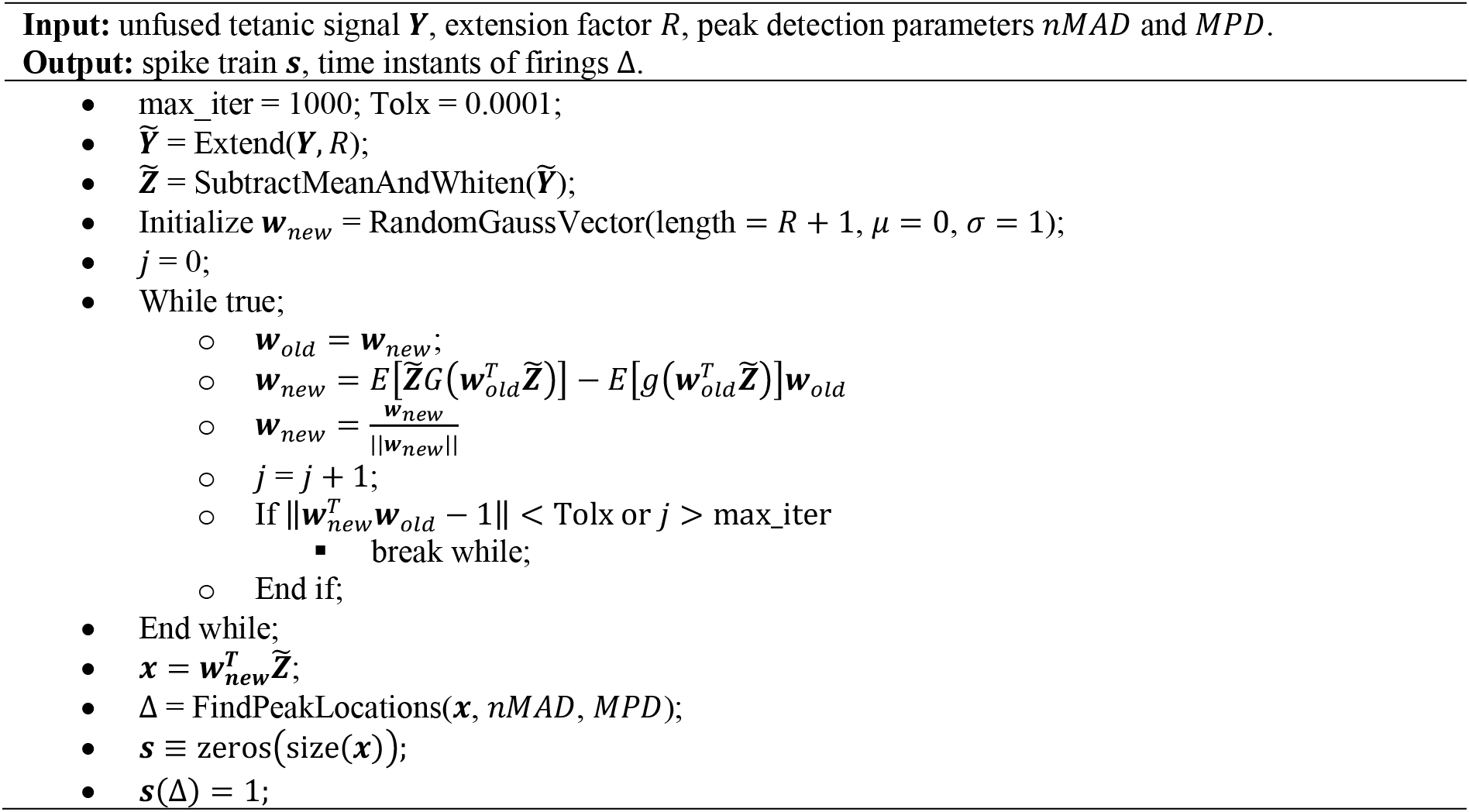

### Parameter and performance evaluation

Based on simulations, we evaluated the algorithms’ three parameters jointly: the extension factor *R* and peak detection parameters *nMAD* and *MPD*. After initial tests to locate the global maxima, we evaluated the following parameters: *R* ∈ [10,15, …,40], *nMAD* ∈ [1.0,1.5, …,4.0], and *MPD* ∈ [10,15, …,40] ms. For each parameter combination, 2,700 unfused tetanic signals were generated based on the combinations of SNR values (30, 20, and 10 dB), ISI CV values (5, 20, and 40%), and firing rates (8, 12, and 16 Hz) as explained in a previous study^17^.

For each generated signal, the rate of agreement (RoA) between the estimated and simulated firings was quantified, defined as 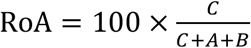, where *C* is the number of correctly identified spikes, *A* is the number of false spikes, and *B* is the number of missed spikes. We set the tolerance interval for correctly identified spikes between 0 and *R* + 10 (extension factor delay) ms, which was selected based on sensitivity analysis (see Fig. S1, Supplementary Material). We also analysed the time between a correctly estimated spike (*C*) and the simulated spike (referred to as spike delta) and its variability (spike delta variability) for different extension factors *R*. The difference in spike delta variation between different extension factors (in parameter evaluation) was tested using Levene’s test^27^ (significance level 5%). The final step of the parameter evaluation was to extract the parameters that maximised the mean RoA (concerning all the simulation parameter combinations).

Using the selected parameters that maximized the mean RoA, the experimental performance of the algorithm was evaluated using two experimental datasets, with the first dataset being the human biceps brachii data and the second dataset being the rat gastrocnemius data. Similar to the parameter evaluation, RoA was calculated for each signal, and we considered the correctly classified spike delta variations.

All data processing and analysis were performed using MATLAB (2022a, MathWorks, Natick, MA, USA).

## Results

### Parameter evaluation – Simulations

The highest mean RoA was for *R* = 20 with 97.5 ± 1.6% (Fig. 4A). Given extension factor *R* = 20, we found that the peak parameters associated with the highest RoA values and smallest standard errors were *nMAD* = 2 and *MPD* = 20 (Fig. 4B-C). However, the latter parameter was not as sensitive as the former.

**Figure 4.**
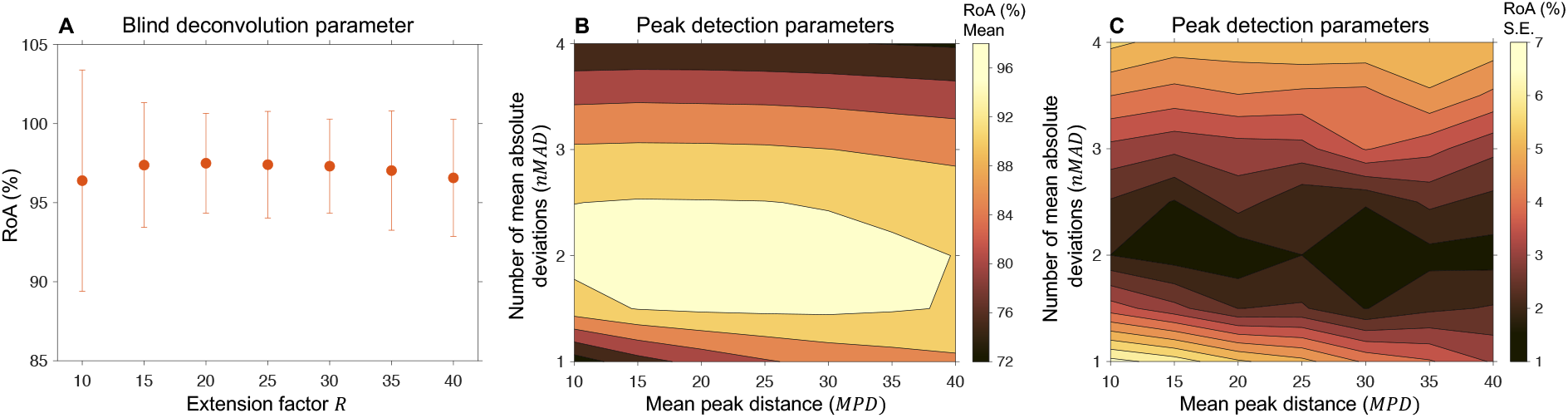
Parameter evaluation of the convolutive blind source separation algorithm using many simulations. **A**. The mean rate of agreement (RoA) and its 95% standard error for different extension factors *R*. Highest mean RoA was for *R* = 20. **B**. Given extension factor *R* = 20, the peak detection parameters (*nMAD* and *MPD*) that had the highest mean RoA were *nMAD* = 2 and *MPD* = 20. **C**. The standard errors associated with the mean RoA values in **B**.

We found that the spike delta differed for different extension factors (Fig. 5A) but also for the same extension factor at different iterations (Fig. 5B). The larger the extension factor, the greater variability in the mean spike delta (Fig. 5C). However, there was no difference (*p* > 0.05) in spike delta variability for the different extension factors (Fig. 5A,B,D). The spike delta variation was 2.2 ± 0.5 ms (Table S2, Supplementary Material).

**Figure 5.**
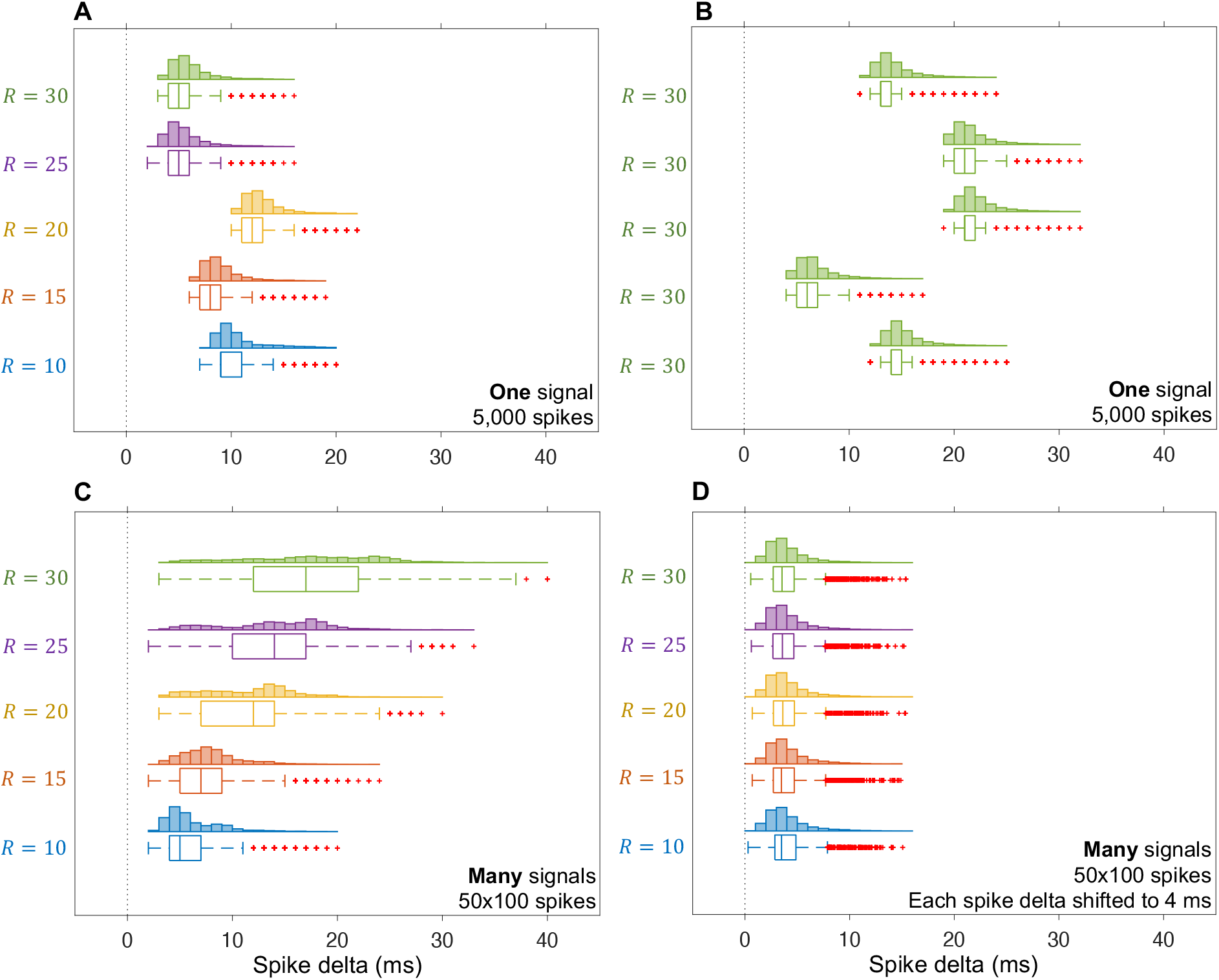
Spike delta and spike delta variability for different extension factors (*R*). **A**. The spike delta differed for different *R*, but there was no difference in spike delta variability. **B**. The spike delta differed for the same *R* for five reiterations of the algorithm, but there was no difference in spike delta variability. **C**. The spike delta differed for different *R*, and there was a difference in spike delta variability. **D**. Subtracting the spike delta for each simulated signal: the spike delta did not differ for different *R*, and there was no difference in spike delta variability.

### Performance evaluation – Experimental data

The three (N=58 spikes) MU velocity-based signals of human biceps brachii had a firing rate of 11.4 ± 2.6 Hz and an ISI CV of 13.4 ± 0.7% (Table 1). The estimated spike trains highly agreed with the EMG reference spike trains (98.0 ± 3.4%) (Table 1). In addition, the spike delta variation was 1.7 ± 0.4 ms.

**Table 1.**
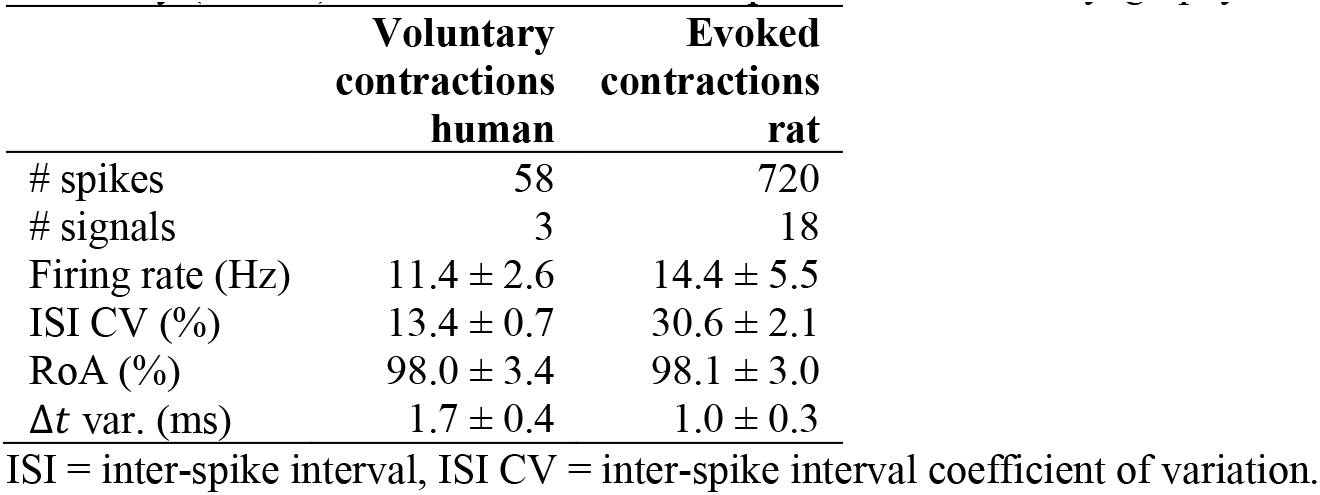
Performance evaluation based experimental signals using the rate of agreement (RoA) and the spike delta variability (Δ*t* var.) between estimated and spikes from electromyography.

The eighteen (N=720 spikes) force-based signals of rat gastrocnemius had a firing rate of 14.4 ± 5.5 Hz and an ISI CV of 30.6 ± 2.1% (Table 1). The estimated spike trains highly agreed with the EMG reference spike trains (98.1 ± 3.0%) (Table 1). In addition, the spike delta variation was 1.0 ± 0.3 ms.

## Discussion

This work aimed to estimate spike trains of simulated and experimental unfused tetanic signals using convolutive BSS. We considered a convolutive BSS algorithm with three parameters based on a fixed-point iteration^10,11^ and peak detection^17^. The main finding was that the estimated spike trains highly agreed with the simulated (97.5 ± 1.6%) and EMG reference spike trains (98.0 ± 3.4% and 98.1 ± 3.0%).

Convolutive BSS of an unfused tetanic signal can be used to estimate its spike train since they highly agree with the simulated (see Fig. 4A) and EMG reference spike trains from two different datasets (see Table 1). We found that the spike delta variability did not depend on the extension factor, although the spike delta differed for different extension factors and the same extension factor (Fig. 5A-B). This observation suggests that there are multiple local maxima, and the convergence to different local maxima may depend on the initialization of the separation vector^10^. Also, a larger extension factor leads to a larger spike delta variance (between trials, Fig. 5C), suggesting that the convergence issue increase with the extension factor value. This problem could be overcome using a more standardized initiation of the separation factor or finding a standardized way to shift the estimated spikes without knowing the ground truth. However, this study aimed to show proof-of-concept of estimating spike trains, and not to optimize the algorithm.

This study considered observations of unfused tetanic signals. Although the results of this study may be used on the estimated unfused tetanic signals from the ultrasound-based pipeline^4,17^, this study provides evidence of including the temporal convolutive BSS approach to improve the separation of displacement velocity images from ultrasound to increase the identification rate. Now we have shown that convolutive BSS can be used to estimate spike trains in the simple case of an unfused tetanic signal. The next challenge emerges when expanding to ultrasound images, i.e., successive twitches within the same MU may be different. There is a possibility that twitches from a MU may be highly similar to the ones of other MUs. However, one could overcome this challenge by including spatial information, which has a high resolution (<1 mm) as the MU relates to a physical component (muscle unit) in the spatial domain. A potential solution may be expanding current sEMG decomposition algorithms^10,11^ to include spatial dependence or sparsity in addition to the temporal deconvolution and validating it using an authentic simulation model^28^. Yet, all these approaches need to deal with the motion of non-MU-related structures that hides a large part of the movement caused by a MU in ultrasound images^12^. One potential solution could be to use the spatiotemporal clutter filtering approach to improve the sensitivity to detect microvascular networks or blood flows corrupted by significant tissue or probe motion artefacts^29^. However, to change the cut-off to increase the sensitivity to MU-associated motion. Nevertheless, the implementation and validation of these approaches will be investigated in the future.

In conclusion, this study estimated spike trains of simulated and experimental unfused tetanic signals using a convolutive BSS algorithm. We found that the estimated spike trains highly agreed with the simulated and EMG reference spike trains. This result implies that the convolutive BSS of an unfused tetanic signal can be used to estimate its spike train. Extending this approach to ultrasound images is promising, but it remains to be investigated in future studies where spatial information is inevitable as a discriminating factor.

## Acknowledgements

We want to thank Professor Jan Celichowski and Rositsa Raikova for providing experimental data on rats irregularly stimulated single motor units. This research received no specific grant.

## References

1. Farina, D., Merletti, R. & Enoka, R. M. The extraction of neural strategies from the surface EMG: an update. J. Appl. Physiol. 117, 1215–1230 (2014).

2. Farina, D., Holobar, A., Merletti, R. & Enoka, R. M. Decoding the neural drive to muscles from the surface electromyogram. Clin. Neurophysiol. 121, 1616–1623 (2010).

3. Rohlén, R., Stålberg, E., Stöverud, K. H., Yu, J. & Grönlund, C. A Method for Identification of Mechanical Response of Motor Units in Skeletal Muscle Voluntary Contractions Using Ultrafast Ultrasound Imaging - Simulations and Experimental Tests. IEEE Access 8, 50299–50311 (2020).

4. Rohlén, R., Stålberg, E. & Grönlund, C. Identification of single motor units in skeletal muscle under low force isometric voluntary contractions using ultrafast ultrasound. Sci. Rep. 10, 1–11 (2020).

5. Ali, H., Umander, J., Rohlén, R. & Grönlund, C. A deep learning pipeline for identification of motor units in musculoskeletal ultrasound. IEEE Access 8, 170595–170608 (2020).

6. Rohlén, R., Yu, J. & Grönlund, C. Comparison of decomposition algorithms for identification of single motor units in ultrafast ultrasound image sequences of low force voluntary skeletal muscle contractions. BMC Res. Notes 15, 207 (2022).

7. Carbonaro, M. et al. Physical and electrophysiological motor unit characteristics are revealed with simultaneous high-density electromyography and ultrafast ultrasound imaging. Sci. Rep. 12, 1–14 (2022).

8. Carbonaro, M., Zaccardi, S., Seoni, S., Meiburger, K. M. & Botter, A. Detecting anatomical characteristics of single motor units by combining high density electromyography and ultrafast ultrasound: a simulation study. in 2022 44th Annual International Conference of the IEEE Engineering in Medicine & Biology Society (EMBC) 748–751 (IEEE, 2022).

9. Holobar, A. & Zazula, D. Multichannel blind source separation using convolution kernel compensation. IEEE Trans. Signal Process. 55, 4487–4496 (2007).

10. Chen, M. & Zhou, P. A Novel Framework Based on FastICA for High Density Surface EMG Decomposition. IEEE Trans. Neural Syst. Rehabil. Eng. 24, 117–127 (2016).

11. Negro, F., Muceli, S., Castronovo, A. M., Holobar, A. & Farina, D. Multi-channel intramuscular and surface EMG decomposition by convolutive blind source separation. J. Neural Eng. 13, 26027 (2016).

12. Lubel, E. et al. Kinematics of individual muscle units in natural contractions measured in vivo using ultrafast ultrasound. J. Neural Eng. (2022). doi:10.1088/1741-2552/ac8c6c

13. Dimigen, O. Optimizing the ICA-based removal of ocular EEG artifacts from free viewing experiments. Neuroimage 207, 116117 (2020).

14. Burke, R. E., Rudomin, P. & Zajac Iii, F. E. The effect of activation history on tension production by individual muscle units. Brain Res. 109, 515–529 (1976).

15. Raikova, R., Celichowski, J., Pogrzebna, M., Aladjov, H. & Krutki, P. Modeling of summation of individual twitches into unfused tetanus for various types of rat motor units. J. Electromyogr. Kinesiol. 17, 121–130 (2007).

16. Raikova, R., Pogrzebna, M., Drzymała, H., Celichowski, J. & Aladjov, H. Variability of successive contractions subtracted from unfused tetanus of fast and slow motor units. J. Electromyogr. Kinesiol. 18, 741–751 (2008).

17. Rohlén, R., Antfolk, C. & Grönlund, C. Optimization and comparison of two methods for spike train estimation in an unfused tetanic contraction of low threshold motor units. bioRxiv (2022).

18. Rohlén, R., Raikova, R., Stålberg, E. & Grönlund, C. Estimation of contractile parameters of successive twitches in unfused tetanic contractions of single motor units – A proof-of-concept study using ultrafast ultrasound imaging in vivo. J. Electromyogr. Kinesiol. 102705 (2022). doi:https://doi.org/10.1016/j.jelekin.2022.102705

19. Fuglevand, A. J., Winter, D. A. & Patla, A. E. Models of recruitment and rate coding organization in motor-unit pools. J. Neurophysiol. 70, 2470–2488 (1993).

20. Milner-Brown, H. S., Stein, R. B. & Yemm, R. Changes in firing rate of human motor units during linearly changing voluntary contractions. J. Physiol. 230, 371 (1973).

21. Tracy, B. L., Maluf, K. S., Stephenson, J. L., Hunter, S. K. & Enoka, R. M. Variability of motor unit discharge and force fluctuations across a range of muscle forces in older adults. Muscle Nerve Off. J. Am. Assoc. Electrodiagn. Med. 32, 533–540 (2005).

22. Lin, D. C., McGowan, C. P., Blum, K. P. & Ting, L. H. Yank: the time derivative of force is an important biomechanical variable in sensorimotor systems. J. Exp. Biol. 222, jeb180414 (2019).

23. Drzymała-Celichowska, H. & Celichowski, J. Functional isolation of single motor units of rat medial gastrocnemius muscle. JoVE (Journal Vis. Exp. e61614 (2020).

24. Krutki, P., Pogrzebna, M., Drzymała, H., Raikova, R. & Celichowski, J. Force generated by fast motor units of the rat medial gastrocnemius muscle during stimulation with pulses at variable intervals. J. Physiol. Pharmacol. an Off. J. Polish Physiol. Soc. 59, 85–100 (2008).

25. Hyvärinen, A., Karhunen, J. & Oja, E. Independent Component Analysis. (Wiley-Interscience, 2001).

26. Hyvärinen, A. Fast and robust fixed-point algorithms for independent component analysis. IEEE Trans. Neural Networks 10, 626–634 (1999).

27. Levene, H. Robust tests for equality of variances. Contrib. to Probab. Stat. Essays Honor Harold Hotell. 279–292 (1960).

28. Ali, H., Umander, J., Rohlén, R., Röhrle, O. & Grönlund, C. Modelling intra-muscular contraction dynamics using in silico to in vivo domain translation. Biomed. Eng. Online 21, 1–19 (2022).

29. Demené, C. et al. Spatiotemporal clutter filtering of ultrafast ultrasound data highly increases Doppler and fUltrasound sensitivity. IEEE Trans. Med. Imaging 34, 2271–2285 (2015).

