## Supplementary Material for "Estimating the neural spike train from an unfused tetanic signal of low threshold motor units using convolutive blind source separation"

### Simulation parameters

Each unfused tetanic signal $Y(t)$ was simulated for the purpose of evaluating the feasibility of using blind deconvolution to estimate the spike train. The simulated data was based on simulating twitches and a spike train.

A twitch model[5] was used for simulating $n$ twitches $F_i, i \in \{1,2,\dots,n\}$ where the analytical equations are presented in Raikova et al. (2008)[5]. The parameters are: the half-contraction time ($T_{hc}$), the contraction time ($T_c$), the half-relaxation time ($T_{hr}$), the twitch duration ($T_{tw}$), and the maximal amplitude ($A_{max}$). The twitch parameters were sampled from a uniform distribution with pre-defined ranges of the parameters suitable for low threshold MUs at low force levels[6] (see Table S1).

The MU firing times $\Delta = (\Delta_1, \Delta_2, \dots, \Delta_n)$, were simulated using a Gaussian renewal process such that $\Delta_{i+1} - \Delta_i \sim N(\mu_{ISI}, \mathrm{CV})$, where $\mu_{ISI}$ and $\mathrm{CV} = \frac{\sigma_{ISI}}{\mu_{ISI}}$ is the mean inter-spike interval (ISI) and coefficient of variation (CV), respectively[1]. To simulate physiological firings and the CV of the ISI at low force levels (low threshold MUs), we chose three firing rates (8, 12, and 16 Hz)[2,3] and three different ISI CVs (5, 20, and 40%)[4].

We summated each twitch to the respective spike resulting in unfused tetanus. Then, the unfused tetanus was differentiated with respect to time. If the twitch is in terms of force, the rate of force development is the yank[8]. If the original twitch is in terms of displacement, the differentiated twitch is in terms of velocity. There are two motivations for using the derivative of unfused tetanus with respect to time:

1) It is commonly stable around its mean value due to the positive and negative parts of the curve approximately cancel each other. Force or displacement twitches are non-stable signals where its shape depends on the MU properties and the instantaneous firing rates.
2) Identifying single muscle units using ultrafast ultrasound is based on decomposing velocity images into spatiotemporal components where the components' temporal information is in terms of (normalized) velocity.

Finally, Gaussian noise was added to the unfused tetanic signal such that the signal-to-noise ratio (SNR) was at a pre-defined level (10, 20, and 30 dB).

### Experimental signals

*Velocity-based unfused tetanic signals from ultrasound on human biceps brachii in vivo*

We retrospectively included, from a previous study, three two-second-long estimated unfused tetanic signals from three healthy subjects (28.3 ± 0.6 years; two females and one male). These signals were a subset of those obtained in a previous study from the cross-section of the biceps brachii at low force levels using the MU ultrasound analysis[3]. The subjects performed low force isometric elbow flexion (90 degrees) and supination of their lower arm as a physician inserted a concentric needle into the biceps brachii muscle and iterated until the physician obtained clear needle EMG signals. The ultrasound probe was positioned proximal to needle insertion such that the image plane captured the needle tip. A concentric needle electrode was used simultaneously with ultrafast ultrasound to record single MU action potentials for each unfused tetanic signal. The ultrasound and EMG systems were synchronized with sampling rates of 2 kHz (radio frequency data of the ultrasound system) and 64 kHz (needle EMG). The needle

EMG data were used to identify the spike train of the MUs as a reference. The subjects gave informed consent before the experimental procedure. The project conformed to the Declaration of Helsinki and was approved by the Swedish Ethical Review Authority (2019-01843). For more detailed information, see Rohlén et al. (2020)[3].

*Force-based unfused tetanic signals from electro-stimulation on functionally isolated motor units in rat gastrocnemius*

We retrospectively included eighteen unfused tetanic signals from a previous study from five adult female Wistar rats. Under pentobarbital anaesthesia, the medial gastrocnemius muscle and the respective branch of the sciatic nerve were partly isolated from surrounding tissues, while other muscles of the hindlimb innervated by the sciatic nerve were denervated. Laminectomy over segments L2-S1 was performed, and the L5 and L4 ventral roots were cut proximally to the spinal cord and split into very thin bundles of axons. The animals were immobilized in a steel frame, and the operated hind limb and the spinal cord were covered with paraffin oil. The Achilles tendon was connected to an inductive force transducer to measure the contractile force under isometric conditions and stretched to the passive force of 100 mN[9]. The thin bundles of axons were electrically stimulated with suprathreshold rectangular pulses (amplitude up to 500 mV, duration 0.1 ms). The all-or-none character of the evoked twitch contractions and action potentials were used as criteria for a single MU isolation[10,11].

The action potentials were recorded with two non-insulated silver wire electrodes (0.15 mm in diameter) inserted through the middle part of the muscle, perpendicular to its long axis. The action potentials were amplified by a multi-channel preamplifier (World Precision Instruments, model ISO-DAM8-A) with a ground electrode inserted into the muscles of the opposite hind limb. The force and EMG signals were sampled at 1 kHz and 10 kHz, respectively.

To ensure that the included unfused tetanic signals were from slow MUs, the MUs were classified based on the shape of 40 Hz unfused regular tetanus. The signals were classified as fast-twitch if a so-called “sag” was observed, otherwise they were classified as slow twitch[12,13]. For three rats, the average stimuli intervals were 100, 80, 70, and 60 ms corresponding to frequencies about 10, 12.5, 14.3, and 16.6 Hz. For two rats, the only the first three stimuli intervals were considered. To mimic voluntary contractions with varying stimuli intervals, the intervals between the individual stimuli were randomly set at values in the mean ISI ± 50% range.

Since we are interested in using the time derivative of the unfused tetanic signal, we wanted to filter the signal before differentiation due to the measurement noise. Each signal was filtered using a $6^{th}$ order zero-phase Butterworth bandpass filter with a high- and low-pass cut-off equal to 3 and 100 Hz. After filtration, the signals were differentiated.

The experimental procedures followed the European Union animal care guidelines, and the principles of Polish Law on the Protection of Animals were approved by the Local Bioethics Committee. For more detailed information, see Drzymała-Celichowska and Celichowski (2020)[14].

### Tables

**Table S1.** The simulation parameters.

| Parameter | Number/Ranges |
|---|---|
| **Unfused tetanus** | |
| Twitches and spikes per simulation | 100 |
| Repeated simulations | 100 |
| SNR (dB) | {10,20,30} |
| **Spike train** | |
| Firing rate (Hz) | {8,12,16} |
| ISI CV (%) | {5,20,40} |
| **Twitch parameters** | |
| $T_{hc}$ (ms) | U[20,25] |
| $T_{c}$ (ms) | U[50,75] |
| $T_{hr}$ (ms) | U[100,130] |
| $T_{tw}$ (ms) | U[300,350] |
| $A_{max}$(n.u.) | U[40,70] |

SNR = signal-to-noise ratio, ISI = inter-spike interval, CV = coefficient of variation.
U[A,B] = the uniform distribution with parameter A and B.

**Table S2.** Parameter evaluation of the convolutive blind source separation algorithm using a large number of simulations and simulation parameters.

| FR (Hz) | ISI CV (%) | SNR (dB) | RoA (%) | Δt (ms) |
|---|---|---|---|---|
| 8 | 5 | 30 | 99.3 ± 0.8 | 0 ± 1.6 |
| 8 | 5 | 20 | 99.7 ± 0.6 | 0 ± 1.9 |
| 8 | 5 | 10 | 99.8 ± 0.5 | 0 ± 2.6 |
| 8 | 20 | 30 | 99.2 ± 0.7 | 0 ± 1.6 |
| 8 | 20 | 20 | 99.4 ± 0.7 | 0 ± 1.9 |
| 8 | 20 | 10 | 98.3 ± 8.7 | 0 ± 2.7 |
| 8 | 40 | 30 | 97.8 ± 1.5 | 0 ± 2.0 |
| 8 | 40 | 20 | 98.4 ± 1.5 | 0 ± 2.3 |
| 8 | 40 | 10 | 96.5 ± 8.8 | 0 ± 2.9 |
| 12 | 5 | 30 | 98.9 ± 0.4 | 0 ± 1.6 |
| 12 | 5 | 20 | 98.9 ± 0.4 | 0 ± 1.8 |
| 12 | 5 | 10 | 98.0 ± 1.2 | 0 ± 2.4 |
| 12 | 20 | 30 | 98.8 ± 0.7 | 0 ± 1.7 |
| 12 | 20 | 20 | 98.8 ± 0.7 | 0 ± 1.9 |
| 12 | 20 | 10 | 97.7 ± 1.4 | 0 ± 2.4 |
| 12 | 40 | 30 | 97.3 ± 1.6 | 0 ± 2.3 |
| 12 | 40 | 20 | 97.4 ± 1.5 | 0 ± 2.4 |
| 12 | 40 | 10 | 94.5 ± 3.2 | 0 ± 3.0 |
| 16 | 5 | 30 | 98.7 ± 0.9 | 0 ± 1.7 |
| 16 | 5 | 20 | 98.2 ± 0.9 | 0 ± 1.9 |
| 16 | 5 | 10 | 94.8 ± 1.9 | 0 ± 2.3 |
| 16 | 20 | 30 | 98.5 ± 0.9 | 0 ± 1.7 |
| 16 | 20 | 20 | 98.1 ± 0.8 | 0 ± 1.9 |
| 16 | 20 | 10 | 93.1 ± 2.6 | 0 ± 2.4 |
| 16 | 40 | 30 | 95.9 ± 2.2 | 0 ± 2.6 |
| 16 | 40 | 20 | 95.2 ± 2.5 | 0 ± 2.9 |
| 16 | 40 | 10 | 88.9 ± 4.1 | 0 ± 3.3 |

FR = firing rate, ISI CV = inter-spike interval coefficient of variation, SNR = signal-to-noise ratio, RoA = rate of agreement, Δt = spike delta.

### Figures

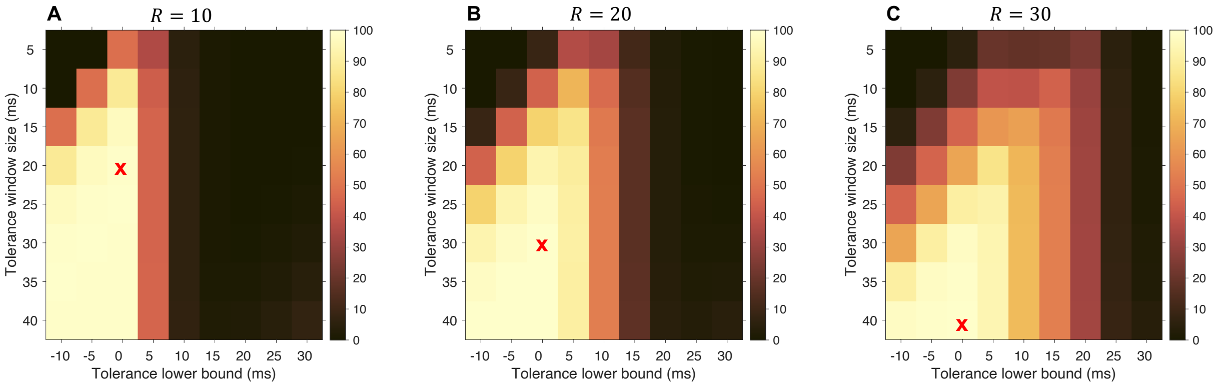

**Figure S1.** Sensitivity analysis of the tolerance interval for rate of agreement (RoA) calculation using different values of extension factor $R$ (10, 20, and 30). We considered a simulation of 5,000 spikes and subsequent twitches using a firing rate of 12 Hz, an ISI CV of 20%, and an SNR of 20 dB. The red crosses denote the selected tolerance intervals.

### References

1. Fuglevand, A. J., Winter, D. A. & Patla, A. E. Models of recruitment and rate coding organization in motor-unit pools. *J. Neurophysiol.* **70**, 2470–2488 (1993).
2. Milner-Brown, H. S., Stein, R. B. & Yemm, R. Changes in firing rate of human motor units during linearly changing voluntary contractions. *J. Physiol.* **230**, 371 (1973).
3. Rohlén, R., Stålberg, E. & Grönlund, C. Identification of single motor units in skeletal muscle under low force isometric voluntary contractions using ultrafast ultrasound. *Sci. Rep.* **10**, 1–11 (2020).
4. Tracy, B. L., Maluf, K. S., Stephenson, J. L., Hunter, S. K. & Enoka, R. M. Variability of motor unit discharge and force fluctuations across a range of muscle forces in older adults. *Muscle Nerve Off. J. Am. Assoc. Electrodiagn. Med.* **32**, 533–540 (2005).
5. Raikova, R., Pogrzebna, M., Drzymała, H., Celichowski, J. & Aladjov, H. Variability of successive contractions subtracted from unfused tetanus of fast and slow motor units. *J. Electromyogr. Kinesiol.* **18**, 741–751 (2008).
6. Rohlén, R. Identification of single motor units in ultrafast ultrasound image sequences of voluntary skeletal muscle contractions. (Umeå University, 2021).
7. Nordez, A. *et al.* Electromechanical delay revisited using very high frame rate ultrasound. *J. Appl. Physiol.* **106**, 1970–1975 (2009).
8. Lin, D. C., McGowan, C. P., Blum, K. P. & Ting, L. H. Yank: the time derivative of force is an important biomechanical variable in sensorimotor systems. *J. Exp. Biol.* **222**, jeb180414 (2019).
9. Celichowski, J. & Grottel, K. The dependence of the twitch course of medial gastrocnemius muscle of the rat and its motor units on stretching of the muscle. *Arch. Ital. Biol.* **130**, 315–324 (1992).
10. Kuffler, S. W., Hunt, C. C. & Quilliam, J. P. Function of medullated small-nerve fibers in mammalian ventral roots: efferent muscle spindle innervation. *J. Neurophysiol.* **14**, 29–54 (1951).
11. Celichowski, J. Motor units of medial gastrocnemius muscle in the rat during the fatigue test. I. Time course of unfused tetanus. *Acta Neurobiol. Exp* **52**, 17–21 (1992).
12. Burke, R. E., Levine, D. N., Tsairis, P. & Zajac Iii, F. E. Physiological types and histochemical profiles in motor units of the cat gastrocnemius. *J. Physiol.* **234**, 723–748 (1973).
13. Grottel, K. & Celichowski, J. Division of motor units in medial gastrocnemius muscle of the rat in the light of variability of their principal properties. *Acta Neurobiol. Exp. (Wars).* **50**, 571–587 (1990).
14. Drzymała-Celichowska, H. & Celichowski, J. Functional isolation of single motor units of rat medial gastrocnemius muscle. *JoVE (Journal Vis. Exp.* e61614 (2020).
